# Dual lineage origins of neocortical astrocytes

**DOI:** 10.1101/2023.09.12.557313

**Authors:** Jiafeng Zhou, Ilaria Vitali, Sergi Roig-Puiggros, Awais Javed, Denis Jabaudon, Christian Mayer, Riccardo Bocchi

## Abstract

Astrocytes represent one of the most abundant cell types in the central nervous system, and play an essential role in nearly all aspects of brain functions^1^. Recent studies have challenged the notion that cortical astrocytes are a uniform population, and have highlighted their diverse characteristics at the morphological, molecular, and functional levels^2-5^. However, how this diversity originates and establishes during cortical development, remains largely unknown. Using single-cell RNA sequencing, we identified five distinct astrocyte subtypes displaying unique spatial patterns in the mouse neocortex, and discovered essential regulators for their formation. Furthermore, we used TrackerSeq^6^, a method that integrates heritable DNA barcodes into the genome of electroporated progenitors, to track clonally related astrocytes, and identified two distinct lineages that give rise to the five astrocyte subtypes. The first lineage derives from Emx1^+^ multipotent progenitors that first generate neurons and then switch to produce cortical astrocytes. The second lineage stems from a fate-restricted progenitor population that exclusively gives rise to a specific subset of cortical astrocytes, marked by Olig2. The knockout of this gene in cortical progenitors is sufficient to promote a fate switch between the two lineages. These findings offer novel insights into the cellular mechanisms underlying astrocyte diversity, highlighting the presence of multiple progenitor subtypes, responsible for generating distinct subtypes of astrocytes.

## Main

Astrocytes are a crucial cell type in the mammalian brain, serving a wide range of physiological functions, including maintenance of neuron viability, formation of the blood brain barrier, and regulation of synapse functions^1^. Although they are generally considered a relatively homogeneous population, recent studies have revealed that astrocyte subtypes displaying distinct molecular and functional features, are present throughout the central nervous system^4,5,7-9^. Additionally, some intra-regional molecular differences have been detected in the neocortex^2,3^. Despite these recent findings, a consensus has still to be reached regarding to which extent cortical astrocytes are heterogenous, and more importantly, the mechanisms by which their diversity is established during development remain largely unknown.

To address these questions, we used a cross-modal approach combining high-throughput single-cell RNA sequencing (scRNA-seq) and Multiplexed Error-Robust Fluorescence in situ Hybridization (MERFISH), that allowed us to identify five molecularly distinct cortical astrocyte subtypes, with specific localizations along the cortical column. Taking advantage of a diffusion-based computational approach (URD)^10^, a method to simulate astrocyte development, we identified key regulators underlying the formation of each subtype. By combining scRNA-seq and massively parallel clonal-tagging (TrackerSeq)^6^ of progenitors, we traced clonally related cortical astrocytes, and provided evidences that astrocyte subtypes arise from two separate lineages, *i*.*e*., progenitors. In addition to the classical multipotent progenitor Emx1^+^, capable to switch its potency to produce first neurons and then astrocytes^11^, we found a fate restricted progenitor population already present at the beginning of the corticogenesis, that gives rise only to a subset of astrocytes expressing the key regulator Olig2. These findings provide an alternative view to the current model, where virtually all astrocytes are generated by multipotent progenitors^11^, transitioning from neurogenesis to gliogenesis^12^. Our results highlight the existence of two distinct progenitor subtypes within the ventricular zone, coexisting and jointly contributing to the generation of cortical astrocytes.

### Molecular identity and spatial distribution reveal five cortical astrocyte subtypes

To investigate the molecular heterogeneity of cortical astrocytes, we produced scRNA-seq datasets from wild type mouse cortices at postnatal day (P)7, a stage when the delamination and expansion of astrocytes are complete^13^. Traditional electroporation of episomal plasmids does not allow permanent labelling of astrocytes, due to their high rate of division and consequent dilution of the plasmid^14^. Therefore, we employed a piggyBac system^15^, utilizing a transposase enzyme (helper plasmid), to integrate the sequence of the green fluorescent protein (GFP) from a donor plasmid into the genome of *in utero* electroporated dorsal progenitors lining the lateral ventricle (Fig. 1a, Extended Data Fig. 1a). To exclude the possibility of losing subpopulations of astrocytes, we targeted progenitors at either embryonic day (E)12.5 or E16.5, and collected all the GFP^+^ cells at P7 using fluorescence-activated cell sorting (FACS) for scRNA-seq (Fig. 1a). These transcriptomic datasets were integrated with a comprehensive atlas of the developing cortex spanning from E10.5 to P4^16^, to facilitate cell type annotations and the identification of the maturation stages (Extended Data Fig. 1b). By employing established gene markers for well-characterized cell populations, we calculated cell-type scores and annotated all known cell types in the cortex, including astrocytes (Extended Data Fig. 1c, d, e). Glial cells, *i*.*e*., astrocyte and oligodendrocyte/oligodendrocyte precursor cells (OPC), were predominantly found in the P7 datasets, while excitatory neurons and progenitors were primarily detected in the embryonic and perinatal datasets (Extended Data Fig. 1f), consistent with the temporal progression of corticogenesis^17^. To specifically investigate astrocyte heterogeneity, we subset the identified astrocytes from the P7 datasets, and re-clustered them for subtype analysis (Fig. 1b). This allowed us to identify five distinct astrocyte subtypes, characterized by unique molecular markers (Fig. 1c, d).

**Fig. 1.**
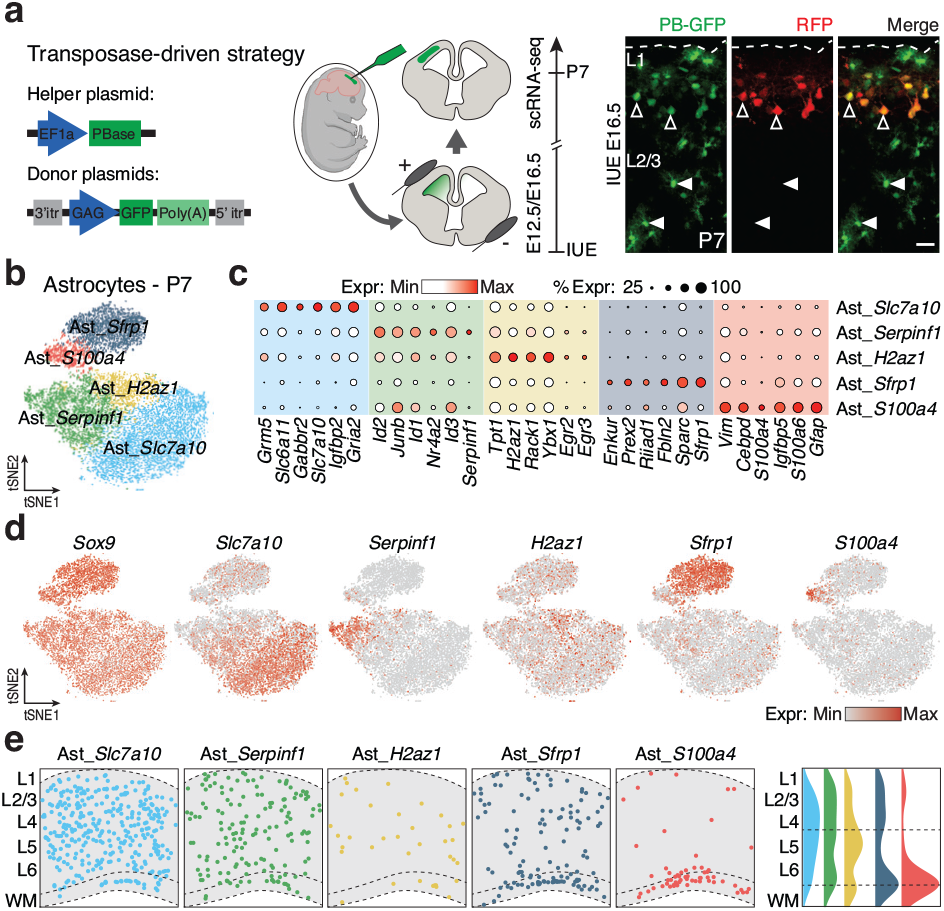
Identification and spatial distribution of astrocyte subtypes in the mouse neocortex. **a**, Left: schematic of the plasmids and of the *in utero* electroporation (IUE) technique employed to label cortical astrocytes. Right: representative confocal images of P7 coronal section electroporated at E16.5 with both integrative (PB-GFP) and episomal (RFP) plasmids. Empty arrowheads indicate GFP^+^/RFP^+^ neurons while full arrowheads indicate GFP^+^/RFP^-^ astrocytes. L1: layer 1; L2/3: layer 2/3. Scale bar: 50 μm. **b**, tSNE plot displaying P7 astrocyte subtypes. **c**, Dot plot showing the expression levels of six representative genes for each of the five identified astrocyte subtypes. **d**, tSNE plot illustrating the expression levels of selected genes for each astrocyte subtypes. *Sox9* gene is a pan-astrocytic marker. **e**, Left: spatial positioning of cortical astrocyte subtypes on an adult MERFISH coronal section. Right: spatial distribution of each astrocyte subtypes along the cortical column.

To predict their cortical location, we took advantage of a publicly available MERFISH spatial dataset^18^, obtained from a coronal section of an adult mouse brain (Extended Data Fig. 1g), which includes a panel of 1122 representative features. Based on these features, we were able to identify all known cell types in the spatial dataset (Extended Data Fig. 1g), subset the astrocyte cluster (Extended Data Fig. 1h, i), and subsequently map the astrocyte subtypes identified in our scRNA-seq datasets, onto the brain section. The five subtypes of cortical astrocytes displayed specific positions along the cortical column (Fig. 1e). For instance, Ast_*Slc7a10*, Ast_*Serpinf1* and Ast_*H2az1* were predominantly enriched in the cortical grey matter (GM) with distinct preferences. In contrast, Ast_*Sfrp1* and Ast_*S100a4* were primarily enriched in the cortical white matter (WM), with Ast_*Sfrp1* also being present in the upper part of the GM (Fig. 1e). These findings support the idea that cortical astrocytes represent a heterogeneous population with five molecularly distinct subtypes, each of them with a specific position within the cortical column.

### Two distinct lineages underlie cortical astrocyte diversity

To gain insights into the establishment of cortical astrocyte heterogeneity during development, we subset and re-clustered progenitor and astrocyte populations from all ages of the integrated scRNA-seq datasets (Fig. 2a and Extended Data Fig. 1c). Based on ages and cell-type scores (Extended Data Fig. 2a, b), we observed a continuum of cell states, transitioning from early progenitors to late progenitors, then to immature astrocytes, and finally to specific astrocyte subtypes (Fig. 2a). Pseudotime analysis revealed a temporal early-to-late gradient, indicating the progression from progenitors to astrocytes, consistent with the expected direction of differentiation^16,19,20^ (Fig. 2b). To investigate the continuum of differentiation during astrogenesis, we used a diffusion-based computational approach (URD)^10^. This approach generated a branched tree structure, based on the transcriptional similarity and pseudotime of cells, providing insights into the specification and differentiation of cortical astrocytes (Fig. 2c, see Methods). We identified a branching point around E16.5, where two major trajectories, named here S100a11 and Olig2, gave rise to the five astrocyte subtypes (Fig. 2c). We then used the reconstructed tree to map transcriptional dynamics along the differentiation trajectories of cortical astrocytes (Fig. 2d, see Methods). Over pseudotime, we observed the downregulation of progenitor genes (*e*.*g*., *Pax6, Sox2, Hes1*), followed by the upregulation of pan-astrocytic genes (*e*.*g*., *Slc1a3, Sox9, Clu*) in both trajectories, albeit with differential temporal patterns. The Olig2 trajectory displayed an earlier downregulation of progenitor genes, along with an earlier expression of pan-astrocytic genes, compared with the S100a11 trajectory. Consistent with this observation, the proportion of astrocyte subtypes across all time points revealed that astrocytes from the Olig2 trajectory emerged at earlier stages (Extended Data Fig. 2c). These evidences suggest that the onset of astrogenesis in the Olig2 trajectory occurs earlier, compared to the S100a11 trajectory. In the later stages of pseudotime, we identified two distinct sets of genes that were specifically enriched in either the S100a11 trajectory (*e*.*g*., *S100a11, Riiad1*, and *Phlda1*), or the Olig2 trajectory (*e*.*g*., *Olig2, Chrdl1*, and *Btbd17*; Fig. 2d, e, Extended Data Fig. 2d). To validate the trajectory analysis, we calculated trajectory scores based on these two gene sets. As expected, the Olig2 trajectory score was higher in Ast_*Slc7a10*, Ast_*Serpinf1* and Ast_*H2az1*, while S100a11 trajectory score demarcated the Ast_*Sfrp1* and Ast_*S100a4* subtypes (Fig. 2f). Together, our molecular analysis identified two primary gene cascades that underlie two distinct trajectories, resulting in the generation of the molecular diversity of cortical astrocytes.

**Fig. 2.**
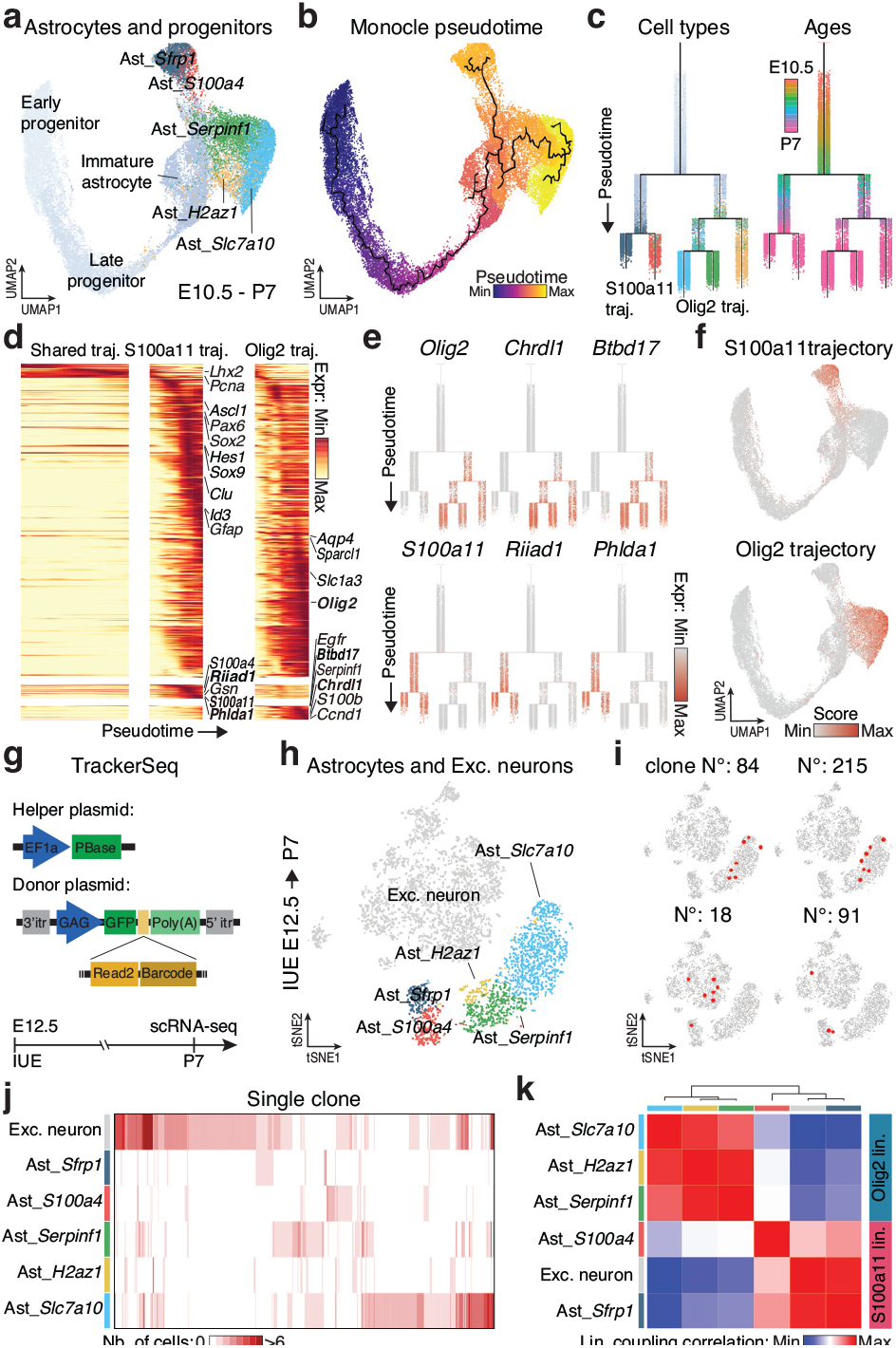
Cortical astrocyte subtypes originate from two molecularly distinct lineages. **a**, UMAP plot displaying progenitors and astrocytes aged from E10.5 to P7. **b**, UMAP plot displaying Monocle pseudotime underlying the developmental trajectory from progenitors to astrocyte subtypes. **c**, URD branching tree simulating the developmental process from progenitors to astrocyte subtypes. Cells are color-coded either by their identities (left) or actual ages (right). **d**, Gene expression cascades of differentially expressed genes between S100a11 and Olig2 trajectories. Genes are ordered based on their temporal dynamics, divided into three segments: shared trajectory (from the top of URD tree to the first branching point of URD tree shown in c), S100a11 trajectory (left clade of URD tree) and Olig2 trajectory (right clade of URD tree). Selected trajectory-specific genes are labelled. **e**, Expression of trajectory-specific genes along URD tree. **f**, UMAP plots depicting S100a11 and Olig2 trajectory scores, calculated based on trajectory-specific genes. **g**, Schematic of the TrackerSeq plasmids and experimental workflow. **h**, tSNE plot of excitatory neurons and astrocyte subtypes traced from E12.5 and collected at P7. **i**, Examples of clones that are either restricted within Olig2 trajectory-derived astrocytes (top) or shared between excitatory neurons and S100a11 trajectory-derived astrocytes (bottom). **j**, Heatmap showing clonal distributions across excitatory neurons and astrocyte subtypes. Each column represents one clone, with number of cells per clone indicated by color scale. **k**, Heatmap of lineage coupling scores between pairs of excitatory neurons and astrocyte subtypes. Values range from positive (red, coupled) to negative (blue, anti-coupled).

Since the clonal relationship between brain cells cannot be accurately inferred by trajectory analyses based on transcriptional profiles alone^6^, we investigated the contribution of clonality to the formation of these trajectories and astrocyte subtypes. To retrieve in parallel molecular identities and clonal relationships of astrocytes, we traced clonally related cortical astrocytes by employing TrackerSeq^6^, a massively parallel tagging system that uses the same piggyBac transposon system mentioned earlier, to permanently insert a GFP reporter gene fused to semi-random synthetic oligonucleotide sequences (barcodes), that can be detected through scRNA-seq (Fig. 2g, top). We co-electroporated the helper and donor plasmids at E12.5, and FAC-Sorted GFP^+^ cells at P7 for scRNA-seq analysis (Fig. 2g, bottom). The resulting dataset included all major cell types of the cerebral cortex, 75% of which were excitatory neurons, astrocytes, and oligodendrocytes/OPC (Extended Data Fig. 3a). For these cell types, we retrieved barcode information from approximately 75% (Extended Data Fig. 3b), allowing us to define clones and examine clonal relationships (see Methods). We first assessed the distribution of clones across the identified cell types (Extended Data Fig. 3c). To reconstruct and quantify clonal relationships among cell types, we proceeded by assessing the probability of recovering shared lineage barcodes from all pairs of cell types, with respect to a random distribution (*i*.*e*., lineage coupling score, see Methods). As expected, hierarchical clustering of the pairwise correlation between coupling scores, revealed a group of lineage-related cell types, including progenitors, neuroblasts and neurons (Extended Data Fig. 3d). Conversely, astrocytes and oligodendrocytes/OPC exhibited a lower coupling correlation with the other cell types, suggesting the existence of glia-enriched lineages (Extended Data Fig. 3d). To obtain a more comprehensive understanding, we recalculated the lineage coupling correlation exclusively amongst subtypes of astrocytes, oligodendrocytes/OPC, and excitatory neurons (Extended Data Fig. 3e). This analysis revelated three main lineages: one corresponding to oligodendrocytes/OPC subtypes, another encompassing a group of astrocyte subtypes only, and a mixed lineage composed of both astrocytes and excitatory neurons. To enhance our understanding of the clonal relationship between astrocyte subtypes and neurons, we subset and re-clustered the astrocyte and neuron clusters from the E12.5-P7 dataset (Fig. 2h). Through the analysis of clone distribution, we found clones that were shared between astrocyte subtypes from the S100a11 trajectory (Ast_*Sfrp1* and Ast_*S100a4*) and excitatory neurons (Fig. 2i, j). The lineage coupling correlation provided further confirmation that astrocytes from S100a11 trajectory exhibited strong connections with excitatory neurons, suggesting a shared developmental origin of these astrocyte subtypes with excitatory neurons (Fig. 2k). Remarkably, we observed the presence of clones exclusively composed of astrocytes from the Olig2 trajectory (Ast_*Slc7a10*, Ast_*Serpinf1* and Ast_*H2az1*; Fig. 2i, j). Indeed, astrocytes belonging to the Olig2 trajectory displayed lower lineage coupling correlation with the neuronal lineage, supporting the notion of a separate origin from excitatory neurons (Fig. 2k). Together our findings unveiled that the five astrocyte subtypes arise from two separate lineages: a multipotent lineage (S100a11 lineage) capable of generating both astrocytes and neurons, and an astrocyte-specific lineage (Olig2 lineage) that exclusively gives rise to astrocytes.

### Two progenitor subtypes account for cortical astrocyte diversity

In order to gain insights into the progenitor subtypes responsible for generating the two lineages, we traced E12.5 progenitors with TrackerSeq, and collected their progenies for scRNA-seq at E18.5, an intermediate timepoint to capture the molecular profiles of various cellular states of progenitors and astrocytes, within a single comprehensive dataset (Fig. 3a, top). As expected, we retrieved different cell types, including progenitors, OPC, excitatory neurons, and the two astrocyte lineages (Fig. 3b, Extended Data Fig. 4a). Using data from Di Bella et al.^16^ as a reference (Fig. 3a, bottom, see methods), we identified two distinct subtypes of progenitors, namely Prog_1 and Prog_2 (Fig. 3b). Moreover, we observed shared clones between Prog_1, S100a11 astrocytes, and excitatory neurons, while Prog_2 clones were predominantly clonally related to the Olig2 lineage astrocytes (Extended Data Fig. 4b). Consistent with these findings, the lineage coupling analysis indicated that Prog_1 was primarily clonally related to the excitatory neurons and astrocytes from S100a11 lineage (Extended Data Fig. 4c). In contrast, Prog_2 showed a stronger clonal relationship with the Olig2 lineage astrocytes. To further enhance the resolution of this analysis, we specifically recalculated the lineage coupling scores only among progenitor subtypes (Prog_1 and Prog_2), excitatory neurons, and astrocyte lineages (Fig. 3c). The analysis confirmed the multipotency of Prog_1, and the fate-restricted capacity of Prog_2, this latter serving as specific origin of astrocyte (Fig. 3c).

**Fig. 3.**
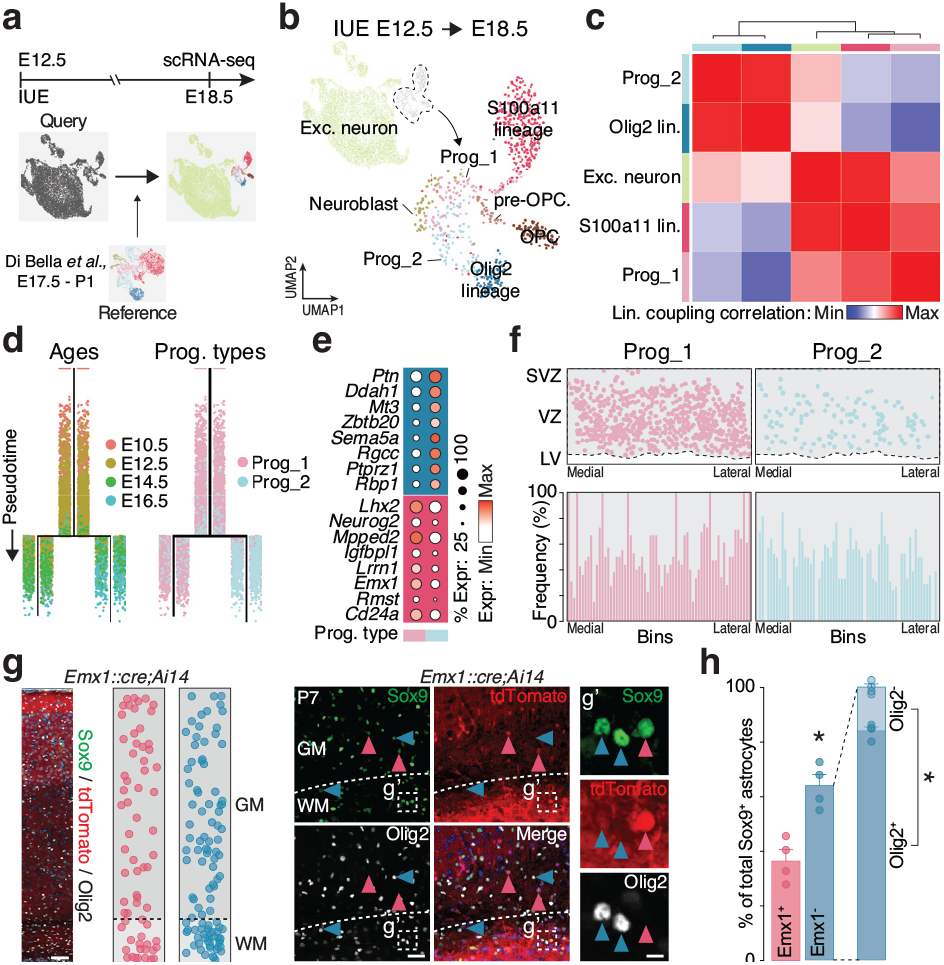
Cortical astrocyte originate from two distinct progenitor subtypes. **a**, Schematic outlining the strategy employed for collecting and annotating E18.5 scRNA-seq dataset. IUE: *in utero* electroporation. **b**, UMAP plot of E18.5 single cells color-coded by cell types. **c**, Heatmap of lineage coupling scores between pairs of E18.5 excitatory neurons, astrocyte lineages and progenitor subtypes. Values range from positive (red, coupled) to negative (blue, anti-coupled). **d**, URD branching trees of progenitors from E10.5 to E16.5, with colors indicating actual ages (left) or predicted identities (right). **e**, Dot plot showing expression of the top 10 genes for each progenitor subtype. **f**, Top: spatial positioning of progenitor subtypes identified from scRNA-seq onto E14.5 MERFISH coronal section. Bottom: distribution of mapped progenitor subtypes along the VZ, divided into 65 bins. LV: lateral ventricle; VZ: ventricular zone; SVZ: subventricular zone. **g**, Left: representative confocal images of a P7 coronal section from *Emx1::cre;Ai14* transgenic mouse immunostained for Sox9 and Olig2. Sox9^+^ astrocytes are divided into two columns based on Emx1 expression (Emx1^+^ and Emx1^-^), each dot represents one astrocyte in the representative image. Right: examples of Sox9^+^ astrocytes: Sox9^+^/Emx1^+^/Olig2^-^ astrocytes are indicated by pink arrowheads while Sox9^+^/Emx1^-^ /Olig2^+^ astrocytes are pointed by blue arrowheads. A magnified view of dashed box (g’) is shown in the rightmost column. Scale bars: 100 μm (left), 50 μm (middle) and 10 μm (right). **h**, Quantifications of Sox9^+^/Emx1^+^ and Sox9^+^/Emx1^-^ astrocytes in the entire cortical column. Olig2 expression is further quantified in all Sox9^+^/Emx1^-^ astrocytes. Values are shown as mean ± s.d.; n = 4 experimental animals; two-tailed *t*-test, \**P* < 0.05.

To explore the heterogeneity of progenitors at early stages, we transferred the cell subtype labels from our E12.5-E18.5 dataset to a reference progenitor dataset spanning E10.5 till E16.5, obtained from the Di Bella et al.^16^ (Extended Data Fig. 4d, top). While most cells were assigned as multipotent Prog_1, we identified a subset of cells labelled as Prog_2 as early as E10.5, indicating the existence of an astrocytic fate-restricted progenitor at the onset of corticogenesis (Extended Data Fig. 4d, bottom). Over time, the number of multipotent Prog_1 decreased, while the population of Prog_2 exhibited an increase from E10.5 to E16.5, suggesting their amplification during this developmental period (Extended Data Fig. 4e, top). To investigate the molecular changes of these two progenitor subtypes throughout corticogenesis, we calculated the average PCA distance within each subtype across different ages (see Methods). We observed that the average PCA distance increased substantially for the multipotent Prog_1 over time, while it remained rather stable for fate-restricted Prog_2. This suggests a temporal cell state progression of Prog_1^21^, giving rise to sequentially distinct neuronal subtypes and later undergoing the neurogenesis-to-gliogenesis switch (Extended Data Fig. 4e, bottom)^12^.

To gain deeper insights into the molecular divergence of Prog_1 and Prog_2 during corticogenesis, we used the URD algorithm to simulate progenitor molecular trajectories with the two progenitor subtype identities at E16.5 as endpoints. The branched structure revealed that the molecular divergence between Prog_1 and Prog_2 starts around E12.5 (Fig. 3d) and is accompanied by the expression of a distinct set of genes, specific to each progenitor subtype (Fig. 3e). Consistent with previous studies, we observed that multipotent Prog_1 highly expressed Emx1, a well-known marker for excitatory neuronal lineage^22,23^. Remarkably, immunostaining on E15.5 *Emx1::cre;Ai14* transgenic mice revealed that only some of Sox2^+^ cortical progenitors expressed Emx1 (Extended Data Fig. 4f), providing evidence that multipotent Prog_1 (Emx1^+^) are not the only ones responsible for astrocyte production.

To further validate the existence of these two progenitor subtypes and predict their spatial localization, we generated a MERFISH spatial transcriptome dataset from a coronal section of an E14.5 mouse brain. We analyzed 464 representative features, which included differentially expressed genes specific to the identified progenitor subtypes (Fig. 3e). Following the annotation of the generated dataset (Extended Data Fig. 5a), we subset the progenitor cluster (Extended Data Fig. 5a, b). We then transferred the identities of the progenitor subtypes Prog_1 and Prog_2 into the spatial progenitor dataset. This analysis unveiled the presence of the two distinct progenitor identities in the ventricular/subventricular zone (Fig. 3f, top). Consistently, we observed previously identified markers, including *Emx1* and *Lhx2*, predominantly expressed by the multipotent Prog_1, while *Zbtb20* and *Ptprz1*, primarily expressed by fate-restricted Prog_2 (Extended Data Fig. 5c, d). Furthermore, we found that Prog_1 and Prog_2 were spatially organized into columnar domains, with each column being formed by alternating between Prog_1 and Prog_2 (Fig. 3f, bottom, Extended Data Fig. 4f), further supporting the co-existence of two distinct progenitor identities.

Finally, to validate these distinct progenitor subtypes and their specific potencies for generating subsets of cortical astrocytes, we crossed *Emx1::cre;Ai14* transgenic mice to specifically trace the S100a11 lineage generated by multipotent Prog_1 Emx1^+^. At P7, we performed immunostaining on coronal brain sections against Sox9 to label all astrocytes, and Olig2 to label astrocytes derived from the Olig2 lineage (Fig. 3g). Remarkably, a small proportion of cortical Sox9^+^ astrocytes derived from Emx1^+^ Prog_1, while a significant majority of Sox9^+^/Emx1^-^ astrocytes were positive for Olig2 (Fig. 3h). These proportions align with those observed in the scRNA-seq (Fig. 1b), MERFISH (Fig. 1e) and most importantly, with the proportion of Olig2^+^ astrocytes determined through immunostaining quantification (see the following paragraph, Fig. 4c, left). Taken together, these findings confirm the co-existence of two distinct progenitor subtypes (multipotent Prog_1 and fate-restricted Prog_2), responsible for generating distinct populations of cortical astrocytes.

**Fig. 4.**
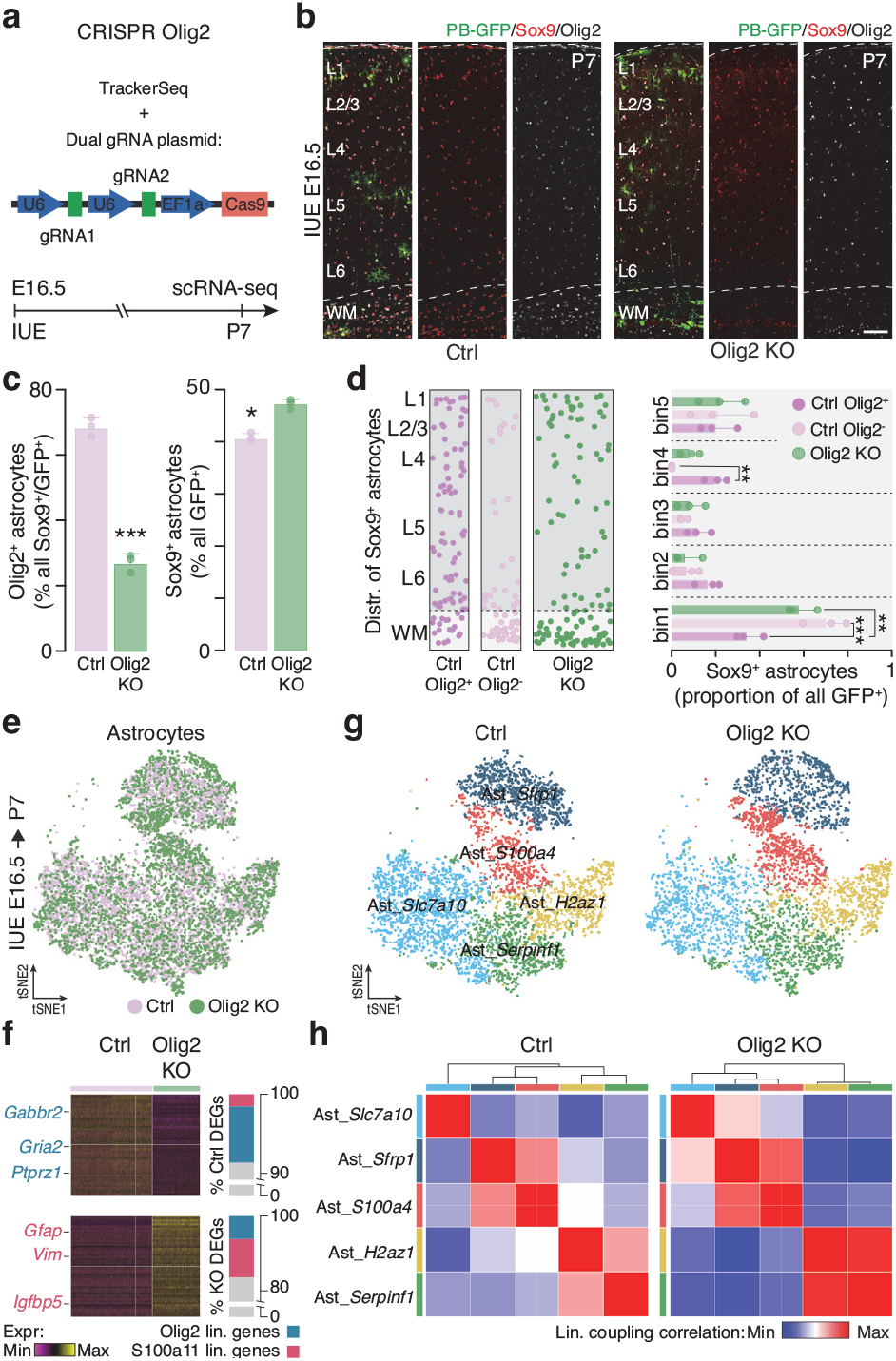
*Olig2* knockout alters the generation of cortical astrocyte lineages. **a**, Schematic of experimental workflow of *Olig2* knockout and lineage tracing. IUE: *in utero* electroporation. **b**, Representative confocal images of cortical columns from control (Ctrl) and *Olig2* knockout (Olig2 KO) brains immunostained for Sox9 and Olig2. Scale bar: 100 μm. **c**, Left: fraction of Olig2^+^ astrocytes in Ctrl and Olig2 KO cortices. Right: fraction of Sox9^+^ astrocytes among all GFP^+^ electroporated cells. Values are shown as mean ± s.d.; n = 3 experimental animals; two-tailed *t*-test, \**P* < 0.05, \*\*\**P*<0.0001. **d**, Left: distribution of Sox9^+^ astrocytes in the cortical column. The control column is separated into two based on Olig2 expression (Ctrl Olig2^+^ and Ctrl Olig2^-^). Each dot represents one cell from all analyzed sections. Right: histograms displaying the distribution of Sox9^+^ astrocytes divided five cortical bins. Values are shown as mean ± s.d.; n = 3 experimental animals; two-way ANOVA with post hoc Tukey test, \*\**P* < 0.01, \*\*\**P*<0.0001. **e**, tSNE plot of astrocytes from both Ctrl and Olig2 KO conditions, traced from E16.5 and collected at P7. **f**, Left: heatmap showing the expression of differentially expressed genes (DEGs) between Ctrl and Olig2 KO astrocytes. Six selected genes are labelled. Right: proportion of lineage-specific genes in common with either Ctrl DEGs or KO DEGs. **g**, tSNE plot of astrocyte subtypes from both Ctrl and Olig2 KO conditions, traced from E16.5 and collected at P7. **h**, Heatmap of lineage coupling scores between pairs of astrocyte subtypes from Ctrl and Olig2 KO conditions. Values range from positive (red, coupled) to negative (blue, anti-coupled).

### Olig2 is the key regulator for fate specification of Olig2 lineage

To provide functional confirmation of the existence of two independent astrocyte lineages, we knocked out *Olig2*, a gene expressed by astrocytes of the Olig2 lineage (Fig. 2d, e). We electroporated TrackerSeq at E16.5, along with a plasmid encoding the Cas9 and two CRISPR gRNAs, to induce the knockout of *Olig2* (Fig. 4a). The Olig2 immunostaining at P7 revealed a significant decrease in the number of Olig2^+^ astrocytes, suggesting that the production of astrocytes from this lineage was disrupted (Fig. 4b, c, left). This decrease was particularly noteworthy considering the slight increase in the overall number of Sox9^+^ astrocytes observed following the Olig2 knockout (Fig. 4c, right). Moreover, we revealed a significant alteration in the spatial distribution of astrocytes after *Olig2* knockout. In control cortices, Olig2 positive astrocytes (Olig2 lineage) distributed more in the GM, while Olig2 negative astrocytes (S100a11 lineage) were enriched in the WM (Fig. 4d left), mirroring the spatial distribution pattern previously revealed by MERFISH (Fig. 1e). Remarkably, under knockout condition, astrocytes exhibited an enrichment in the WM, similar to the distribution of Olig2 negative astrocytes in control condition (Fig. 4d left). To quantify this observation and compare the astrocyte allocation, we measured astrocyte distribution across 5 cortical bins, which evenly divide the whole cortical column. Compared to the distribution of Olig2 positive astrocytes under control condition, a significantly higher number of Sox9^+^ astrocytes occupied the WM in knockout cortex, mirroring the distribution of Olig2 negative astrocytes in control cortex (Fig. 4d, right). The similar spatial distribution between *Olig2* knockout astrocytes and Olig2 negative astrocytes in control condition suggests a convergence towards a more similar molecular identity after knockout.

To assess whether the loss of *Olig2* led to a reassignment in the molecular identity of the affected cells, we performed scRNA-seq analysis on FAC-Sorted GFP^+^ cells at P7 from both the control and *Olig2* knockout brains electroporated at E16.5 (Fig. 4e). Differential gene expression analysis between these two conditions revealed the expected downregulation of genes typically expressed by astrocytes belonging to the Olig2 lineage (Fig. 4f, Extended Data Fig. 6a). Surprisingly, alongside this decrease, we observed an upregulation of genes typically expressed by the S100a11 lineage, suggesting a fate switch following *Olig2* knockout (Fig. 4f, Extended Data Fig. 6a). To confirm this hypothesis, we first annotated astrocytes into the five different subtypes previously identified (Fig. 4g, see Methods). In line with the immunostaining analysis (Fig. 4b, c), we observed an overall decrease in the number of Olig2 lineage astrocytes, along with an increase in astrocytes belonging to the S100a11 lineage (Extended Data Fig. 6b). Interestingly, after *Olig2* knockout, the lineage coupling analysis revealed a closer clonal relationship between the two astrocyte lineages compared to control condition (Fig. 4h), with an increased number of shared clones between the Olig2 and S100a11 lineages (Extended Data Fig. 6c). The alteration in lineage relationship fosters Olig2 as a necessary key regulator to specify progenitors towards the Olig2 lineage, therefore allowing a preferential differentiation into astrocytes of the S100a11 lineage upon the knockout of *Olig2*. Taken together, these results support the idea that two molecularly distinct progenitors and their respective lineages contribute to the formation of cortical astrocytes heterogeneity.

## Discussion

Our data provide a comprehensive new perspective on how astrocyte heterogeneity forms during the development of the mouse neocortex. We highlighted the existence of two distinct progenitor subtypes, the classical multipotent progenitor Emx1^+^, capable to switch its potency to produce first neurons and then astrocytes^11,12^, and a fate restricted progenitor, giving rise to a specific subset of astrocytes, both coexisting and jointly contributing to the generation of all dorsally derived cortical astrocytes.

Through the combination of *in utero* electroporation and the piggyBac transposase system, we have successfully achieved a permanent labeling of cortical astrocytes, enabling us to exclusively focus our study on astrocytes originating from the dorsal pallium. Using this approach, we first revealed that mouse cortical astrocytes can be classified into five molecularly distinct subtypes. Including WM astrocytes in our analyses, provided us an opportunity to explore the astrocyte diversity also within this region, which has been barely considered in previous studies^2,3,5^. While the most significant molecular disparities were observed among astrocytes residing in either the GM or WM, our analysis also revealed diversities within these two regions, further confirming the recent notion that cortical astrocytes constitute a diverse population^2-5,7-9,24^. Moreover, we identified variations in their spatial distribution within the cortical column. Some subtypes were predominantly localized at specific levels of the cortical column, while others exhibited a more widespread presence. These distinctions in spatial distribution, along with their molecular differences, suggest that certain astrocyte subtypes primarily found in specific locations may have related specialized functions (*e*.*g*., tuning different neural circuits)^25^. In contrast, others may be associated with executing common functions (*e*.*g*., brain energy metabolism)^26,27^, explaining their ubiquitous distribution across the entire neocortex.

Nevertheless, the precise contributions of intrinsic and extrinsic mechanisms to the development of cortical astrocyte heterogeneity remain poorly understood, and a consensus regarding the establishment of this diversity during cortical development has yet to emerge. A combinatorial fluorescence-based labeling strategy was recently used to claim that astrocyte colonization of the neocortex occurs in what appears to be a random manner, rather than being guided by intrinsic and clonally related programs^13^. Furthermore, the proper formation of astrocyte subtypes and their spatial organization also seems to be also influenced by neuronal layers^2,3^. Recent research has revealed that cortical lamination mutants, such as *Reeler* and *Satb2-KO* mice, also exhibit altered glial layering and subtype formations. In contrast, using a similar combinatorial labeling strategy, another study highlighted that protoplasmic astrocytes populating the GM derive from largely separated lineages as opposed to pial astrocytes, supporting the notion that cell intrinsic factors could contribute to drive astrocyte heterogeneity, at least in part^28^. Our study harnesses the power of TrackerSeq^6^, an electroporation-based technique that unbiasedly labels progenitors, to concurrently uncover clonal relationships (*i*.*e*., origins) and transcriptomic signatures (*i*.*e*., identities) at single cell level. This approach unveiled, for the first time, the critical influence of clonality *i*.*e*., cell origins in molding the distinct identities of cortical astrocytes.

We have unveiled two distinct progenitor subtypes responsible for the emergence of cortical astrocytes. A subset of astrocytes is derived from the classical multipotent Emx1^+^ progenitor, capable of generating both excitatory neurons and astrocytes. This discovery is consistent with the outcomes of the MADM lineage tracing experiment conducted using *Emx1::CreER*^*T2*^ mice, where roughly 16% of neuronal clones were identified to contain both neurons and astrocytes^11^. Furthermore, both studies provide corroborative evidence that all neuronal subtypes originate from common multipotent Emx1^+^ progenitors^11^. Remarkably, we revealed that astrocytes generated by multipotent Emx1^+^ progenitors represent a minority of all cortical astrocytes, while the majority of cortical astrocytes originate from fate-restricted astrocytic progenitors already present at the onset of corticogenesis (*i*.*e*., E10.5). This discrepancy with the MADM results can be attributed to the fact that in the above-mentioned study, *Emx1::CreER*^*T2*^ was used to activate the MADM system, which inadvertently excluded fate-restricted astrocytic progenitors that do not express Emx1^11^. Moreover, we found that astrocytes originating from fate-restricted progenitors emerge earlier during corticogenesis, than those born from the shared neuronal lineage. This is in line with a previous study that demonstrated astrocyte progenitors at E17-E18 predominantly contributing to the production of GM astrocytes^29^, which we described as primarily originating from fate-restricted progenitors. The delayed production of astrocyte in S100a11 lineages might be explained by the fact that the Emx1^+^ progenitors must undergo a change in their competencies to allow the neurogenesis-to-gliogenesis switch to happen, once the neurogenesis is completed. Finally, the existence of these two separate progenitor populations is further supported by their spatial organization in columnar domains. This can be facilitated by several families of cell surface receptors, providing homophilic (*e*.*g*., cadherins) or heterophilic (*e*.*g*., integrins) cell-cell adhesive interactions^30,31^. Notably, we also found several cadherins (*i*.*e*., *Cdh4*) differentially expressed between the two subtypes of progenitors, which might explain this columnar organization.

Finally, we provided evidences that *Olig2* plays a critical role in regulating the competences of these fate-restricted subset of progenitors. The knockout of *Olig2* resulted in a reduction of Olig2^+^ astrocytes, accompanied by a shift in their identity toward the S100a11 lineage. Consistent with previous observations^32^, we observed an increase in GFAP expression, a marker highly expressed by the S100a11 lineage. These findings collectively emphasize the substantial influence of cell-intrinsic genes and clonality in determining astrocyte identities.

Taken together, our data show that cortical astrocyte subtypes possess distinct molecular and spatial features and, more importantly, stem from two subtypes of progenitors that coexist throughout corticogenesis. Gaining a deeper understanding of the lineage relationships and origins of cortical astrocytes has the potential to uncover crucial astrocyte subtype-specific mechanisms underlying astrocyte-related developmental disorders, including RASopathies and autism spectrum disorders^33^. Furthermore, in the last decade, astrocytes have emerged as highly promising candidates for targeted neuronal reprogramming^34,35^. The possibility that specific astrocyte subtypes which share their origin with excitatory neurons, like the S100a11 lineage astrocytes, could be more susceptible to reprogramming into neurons, remains to be tested. This potential susceptibility presents a unique opportunity to enhance and refine cell therapy approaches for brain repair, potentially increasing their effectiveness.

## Methods

### Mouse strain

Female mice of the CD1 and C57BL/6 strains, obtained from Charles River Laboratory, were utilized in this study. The embryos were staged based on days post-coitus, with E0.5 defined as 12:00 on the day following the detection of a vaginal plug after overnight mating. For lineage tracing of Emx1^+^ progenitors, Emx1-IRES-Cre mice (B6.129S2-Emx1^tm1(cre)Krj/^J, #005628) were crossed with Ai14 mice (B6.Cg-Gt(ROSA)26Sor^tm14(CAG-tdTomato)Hze^/J, #007914), both obtained from Jackson Laboratory. The experimental procedures detailed in this study were conducted in strict adherence to Swiss laws and had been previously approved by the Geneva Cantonal Veterinary Authority. All mice were housed in the institutional animal facility, with a standard 12-hour light and 12-hour dark cycle and had access to food and water ad libitum.

### In utero electroporation

In utero electroporation was conducted as previously described^36^. Briefly, timed pregnant CD1 mice were anesthetized using isoflurane (5% for induction, followed by 2.5% during the surgical procedure), and they received the analgesic Temgesic from Reckitt Benckiser (10% in 0.9% NaCl). Embryos were injected in lateral ventricle with ∼1 μl of DNA plasmid solution (diluted in endotoxin-free water and 0.002% Fast Green FCF (Sigma)). For the TrackerSeq experiments, the plasmid solution comprised 0.5 ng/μl of Helper plasmid and 0.5 ng/μl of Donor plasmid^6^. In the case of the Olig2 knockout experiments, the plasmid solution consisted of 0.5 ng/μl of Helper plasmid, 0.5 ng/μl of Donor plasmid, and 0.5 ng/μl of the Olig2 CRISPR plasmid (pX458-Ef1a-Cas9-H2B-mCherry, Addgene #171101). The electroporation of embryos was achieved by securing each head between circular tweezer-electrodes (5 mm in diameter, Sonidel), positioned across the uterine wall. Subsequently, a square-wave electroporator (Nepa Gene, Sonidel) delivered either 5 electric pulses at 25 V for 50 ms at 1 Hz for E12.5 embryos, or 5 electric pulses at 45 V for 50 ms at 1 Hz for E16.5 embryos.

### Immunohistochemistry

For tissue preparation, postnatal mice underwent cardiac perfusion with a 4% PFA solution, after which their brains were fixed by immersion in 4% PFA at 4°C overnight, and subsequently stored in PBS at 4°C. Coronal brain sections of 70 *μ*m in thickness were generated using a vibrating microtome (Leica, VT1000S). In the case of embryonic brains, they were fixed for a duration of 4 hours using 4% PFA, cryoprotected by immersion in 20% sucrose-PBS at 4°C overnight. They were then embedded in O.C.T. (CellPath, KMA-0100-00A), snap-frozen at -70°C in isopentane and stored at -80°C. Coronal sections with a thickness of 16 *μ*m were produced employing a cryostat (Leica, CM3050). For the staining process, both types of sections were initially subjected to a 1-hour pre-incubation at room temperature within a solution that simultaneously blocked and permeabilized, consisting of 5% bovine serum albumin and 0.3% Triton X-100 in PBS. Subsequently, they were incubated with primary antibodies overnight at 4°C. Afterward, the sections underwent three rinses in PBS and were then exposed to corresponding Alexa-conjugated secondary antibodies (1:1000; Invitrogen; Donkey anti-Rabbit IgG, Alexa Fluor 488 A11055; Donkey anti-Rabbit IgG, Alexa Fluor 647 A31573; Donkey anti-Goat IgG, Alexa Fluor 488 A21206) for 2 hours at room temperature. Finally, the sections were mounted using Fluoroshield with DAPI (Sigma-Aldrich, F6057). The primary antibodies used, along with their dilutions, were as follows: rabbit anti-Sox9 (Merck Millipore, AB5535, 1:500), goat anti-Olig2 (R&D Systems, AF2418, 1:200), rabbit anti-Sox2 (Abcam, ab97959, 1:500). The images were captured using a ZEISS LSM 800 Airyscan confocal microscope.

### TrackerSeq library preparation

TrackerSeq represents a piggyBac transposon-based library designed explicitly for seamless integration with the 10x single-cell transcriptomic platform. The creation of the TrackerSeq library followed a previously established procedure^6^. In summary, the piggyBac donor plasmid (PBCAG-eGFP, Addgene #40973) underwent several modifications. First, a Read2 partial primer sequence was inserted into the 3′ UTR of the eGFP sequence. Subsequently, through six distinct Gibson Assembly reactions (NEB, #E2611S)^37^, the synthetic lineage barcode oligo mix was cloned between the Read2 partial primer and the poly(A) tail of eGFP. These modifications enable the retrieval of barcodes by the 10x single-cell transcriptomic platform.

### Sample collection for scRNA-seq

Brains collected from E18.5 embryos were dissected on ice with Leibowitz medium with 5% FBS whereas brains collected from P7 pups were exposed and kept in bubbled EBSS with 5% FBS on ice then transferred to Hibernate A medium with 10% FBS and B27 (1:50 dilution), while being observed under a dissecting scope to identify the positive regions. GFP^+^ cortices were then dissociated using the Papain dissociation system following the recommended protocol from Worthington (#LK003150), and further processed with the gentleMACS Dissociator following the manufacturer’s instructions. To isolate and collect GFP^+^ cells, flow cytometry was employed with the Beckman Coulter Moflo Astrios FAC-sorter. Initially, cell suspensions were gated based on DAPI-negative and forward scatter, and from within this population, cells expressing GFP were collected in bulk for subsequent processing on the 10x Genomics Chromium platform. In all FACS experiments, non-electroporated brain tissue served as a negative control to exclude background fluorescence.

### Construction of scRNA-seq and TrackerSeq libraries

For experiments involving the 10x Genomics platform, the following materials were employed: Chromium Single Cell 3′ Library & Gel Bead Kit v3.1 (PN-1000121), Chromium Single Cell 3′ Chip Kit v3.1 (PN-1000127), and Dual Index Kit TT Set A (PN-1000215). These reagents were utilized in accordance with the manufacturer’s provided instructions. The lineage/TrackerSeq barcode library amplification process followed the previously outlined procedure^6^, using the standard NEB protocol for Q5 Hot Start High-Fidelity 2X Master Mix (#M094S) in a 50-μl reaction, with 10 μl of cDNA as the template. To be specific, each PCR reaction contained the following components: 25 μl Q5 High-fidelity 2X Master Mix, 2.5 μl of a 10 μM P7_indexed reverse primer, 2.5 μl of a 10 μM i5_indexed forward primer, 10 μl of molecular-grade H20, and 10 μl of cDNA.

### Processing of sequencing reads

Transcriptome libraries and TrackerSeq barcode libraries were subjected to sequencing using the NovaSeq 6000 system from Illumina, conducted at the iGE3 genomic platform at the University of Geneva and MPIB Next Generation Sequencing core facility. The sequencing data in FASTQ files were subsequently aligned to a reference transcriptome (mm10-2.1.0) and collapsed into Unique Molecular Identifier (UMI) counts using the 10x Genomics Cell Ranger software, versions 6.0.1. In the case of TrackerSeq, preprocessing of the reads in the R2 FASTQ files involved trimming the sequences situated to the left and right of the lineage barcodes (BC). Lineage barcodes shorter than 37 base pairs were excluded from further analysis. Cell barcodes (Cell) were extracted from the corresponding Seurat dataset to create a whitelist of cell barcodes. These extracted cell barcodes and UMIs were incorporated into the read names of the lineage barcode FASTQ files. The resulting FASTQ files underwent processing to generate a sparse matrix in CSV format. In this matrix, the rows represented individual cells identified by their unique cell barcodes, while the columns represented lineage barcodes. Only Cell–UMI–BC triples supported by a minimum of 10 reads and Cell–BC pairs with at least 6 UMIs were considered for subsequent analyses. The assignment of CloneIDs to cell barcodes was accomplished by clustering the matrix using Jaccard similarity and average linkage, following the methodology demonstrated by Wagner and colleagues^6,38^.

### Lineage coupling z-scores and correlations analysis

We conducted an analysis to determine shared clones, lineage coupling z-scores, and correlations for each pair of cell types or cell subtypes, following previously established methods^6,38^. Briefly, a TrackerSeq clone was deemed “shared” if it included a minimum of 2 individual cells assigned to each state. We tallied the total count of shared TrackerSeq clones for each pair of cell types or cell subtypes. This count was then compared to randomized data in which assignments of cell types or cell subtypes were randomly shuffled. We performed 10,000 random permutations to calculate a z-score for each original count of “shared” clones in relation to the expected distribution by random chance. The lineage coupling z-scores were then subjected to hierarchical clustering and visualized as a heatmap. Positive lineage coupling z-scores indicated pairs of cell types or cell subtypes that shared significantly more TrackerSeq clone barcode hits, than what would be expected by chance, while negative lineage coupling z-scores suggested significantly less coupling than expected by chance. Furthermore, we computed correlation coefficients between the z-scores for each pair of cell types or cell subtypes and represented these lineage coupling correlations as a clustered heatmap. For a more detailed explanation of the calculation of lineage coupling z-scores and correlations, please refer to the original TrackerSeq study^6^.

### Data integration, annotation and subtype identification

All scRNA-seq analysis were performed in R (version 4.2.1) following the Seurat workflow (version 4.3.0.1)^39^ for cell filtering and data normalization. Each dataset was initially imported into R as a count matrix, which was then transformed into a Seurat object using the CreateSeuratObject() function with the following parameters: min.cells=3 and min.features=200. Cells harboring more than 10% mitochondrial genes were excluded from further analysis. To integrate the P7 datasets with those from Di Bella et al.^16^ (Extended Data Fig. 1c), we employed Harmony (version 0.1.1)^40^ using the RunHarmony() function with default parameters. For the identification of Seurat clusters, we constructed a shared-nearest neighbor graph using the FindNeighbors() function, based on Harmony embeddings (dimensions = 30). This graph served as input for the SLM algorithm, implemented through the FindClusters() function in Seurat (dimensions = 30, res = 0.5). Concurrently, cell type scores were computed using known brain cell type markers via the CellCycleScoring() function. Clusters were manually annotated based on these cell type scores, gene expression, as well as using the online available databases for the mouse brain (http://mousebrain.org)^41^. Clusters that could not be definitively assigned to a specific cell type were labeled as “Undefined.”

To identify astrocyte subtypes at P7 and distinguish them from more immature states, we initially isolated all astrocytes and progenitors from the integrated dataset (Extended Data Fig. 1c). We then conducted a re-clustering step using Seurat functions FindNeighbors() and FindClusters() with dimensionality set to 40 and a resolution of 1.3 (Fig. 2a). Astrocyte clusters were named based on top regulated genes and their annotation was kept throughout all the datasets of this study (Fig. 1b, Fig. 4g).

To identify progenitor subtype- and astrocyte-lineages at E18.5 (Fig. 3b), we began by manually annotating the E18.5 dataset into general cell types (e.g., excitatory neurons, astrocytes, progenitors) based on Seurat clusters and cell type scores. To facilitate the annotation of cell subtypes, including progenitors and astrocyte lineages, we generated a reference dataset sourced from Di Bella et al.^16^, which encompassed progenitors and glia cells across E17.5 to P1 timepoints.

For predicting progenitor subtypes before E16.5 (Extended Data Fig. 4d), we employed the FindTransferAnchors() and TransferData() functions to transfer labels from the E18.5 dataset to a progenitor dataset derived from the Atlas of Di Bella et al.^16^. This progenitor dataset was created by isolating progenitors from E10.5, E12.5, E14.5, and E16.5 time points (E10.5-E16.5 dataset).

To identify astrocyte subtypes within the Olig2 KO dataset, we integrated both the WT and Olig2 KO datasets using the Harmony function RunHarmony() with default parameters. Subsequently, we subset the astrocyte population and conducted a re-clustering step using the Seurat functions FindNeighbors() and FindClusters(). The dimensionality was set to 30, and the resolution was set at 0.8. Given the substantial overlap between the WT and Olig2 KO astrocytes, and the presence of subtype labels for the astrocytes from the WT dataset (Fig. 1b), we assigned the same labels to astrocytes from the Olig2 KO dataset that fell within the same cluster (Fig. 4g).

### Differential gene expression analysis

To identify markers specific to astrocyte subtypes (Fig. 1c), we used the Seurat function FindAllMarkers() with the following parameters: min.pct = 0.25 and logfc.threshold = 0.25. For the identification of markers associated with progenitor subtypes (Prog_1 and Prog_2, Fig. 3e), we used the function FindMarkers() with parameters set to min.pct = 0.2 and logfc.threshold = 0.2. To identify genes that were differentially expressed between the Ctrl and Olig2 KO datasets (Fig. 4f), we utilized the Seurat function FindAllMarkers() with the following parameters: min.pct = 0.25, logfc.threshold = 0.25, and only.pos=T. Genes with an adjusted p-value greater than 0.05 were retained for subsequent analysis.

### MERFISH analysis

MERFISH experiment was performed following instructions from VIZGEN MERSCOPE Platform. For tissue processing and sectioning: embryos were collected at E14.5, in cold HBSS. The obtained tissue was rapidly fresh-frozen in cold isopentane and preserved at -80°C. We performed 10µm coronal sections using a Leyca CM3050 cryostat and collected them on MERSCOPE slides. Briefly, sections were postfixed for 10 minutes in fresh 4% PFA and preserved for 24h in 70% EtOH. This was followed by the 464 gene panel incubation o/n at 37°C. After the hybridization, the section was embedded in a gel solution and cleared. Finally, the slide was placed in the MERSCOPE instrument to run the acquisition pipeline (https://vizgen.com/). Gene expression matrix and coordinates were exported from the MERSCOPE Visualizer for each ROI and assembled in a Seurat project for gene expression analysis. Briefly, we manually defined ROIs (each one corresponding to a single cell) using the MERSCOPE visualizer (https://vizgen.com/vizualizer-software/). We then exported the raw gene counts per ROI (for the 457 detected genes) and processed single-cell expression following the Seurat workflow (version 4.3.0.1): (1) gene count normalization to the total expression and log-transformation, (2) highly variable genes detection and principal component analysis, (3) graph-based clustering (with the 30 first principal components and a clustering resolution of 0.8), (4) tSNE calculation. For cell type annotation, we calculated cell type scores for both adult and E14.5 MERFISH datasets and manually annotated Seurat clusters into cell types. To predict the position of astrocytes in the adult cortex, we used our P7 astrocyte dataset (Fig. 1b) as the reference, and transferred the astrocyte subtype labels to the adult MERFISH dataset, via the functions FindTransferAnchors() and TransferData() from Seurat. For the prediction of the position of progenitor subtypes in E14.5 MERFISH dataset, same functions were used with the E10.5-E16.5 dataset (Extended data Fig. 4d) used as reference.

### Inference of developmental trajectories of astrocytes

To reconstruct the transcriptional trajectories underlying the specification and differentiation processes of the astrocyte population, URD^10^ was employed. The analysis began by subsetting all progenitors and astrocytes from the integrated dataset (Fig. 2a), serving as the initial dataset for URD simulation. First, we calculated a diffusion map using Destiny v2.14.044 implemented in the function calcDM() from URD with knn = 200. As root cells, we choose a subset of progenitors at E10.5. Cells were ordered in accordance with pseudotime by simulating diffusion from the root and subsequently calculating the distance of each cell from the root. For this, we used the function floodPseudotime() with parameters: n = 100 (number of simulations) and minimum.cells.flooded = 2, followed by the functions floodPseudotimeProcess() and pseudotimeDetermineLogistic(). We further used pseudotimeWeightTransitionMatrix() with default parameters to determine the slope and inflection point of the logistic function used to bias the transition probabilities. As tip cells, we assigned P7 astrocyte subtypes, based on which we simulated random walks on the cell-cell graph from each tip to the root using connections in the biased transition matrix via the functions simulateRandomWalk() and processRandomWalks() from URD. Specifically, 12,500 random walks were performed for each tip. Finally, the URD tree was built using function buildTree() with following parameters: visit.threshold = 0, minimum.visits = 0, bins.per. pseudotime.window = 2, cells.per.pseudotime.bin = 300, divergence. method = “ks”, p.thresh = 0.025, dendro.node.size = 150, min.cells.per.segment = 500 and min.pseudotime.per.segment = 0.01.

### Gene cascade analysis

To identify marker genes specific for trajectories, we used the function aucprTestAlongTree() in the URD package to work backward from each tip along the trajectory, making pairwise comparisons between the cells in each segment and the cells from each of that segment’s sibling and children (segments with equivalent or higher pseudotime values)^10,16^, with following parameters: log.effect.size = 0.25, auc.factor = 1, max.auc.threshold = 1, frac.must.express = 0.1, frac.min.diff = 0.1, genes.use = genes.use, root = 10, only.return.global = False and must.beat.sibs = 0.5. We also used the funcion markersAUCPR() to calculate marker genes for each tip with following parameters: effect.size = 0.4 and auc.factor = 1.1. We than combined marker genes identified from functions aucprTestAlongTree() and markersAUCPR(). Marker genes for Ast_*Slc7a10*, Ast_*Serpinf1* and Ast_*H2az1* were grouped as Olig2 trajectory marker genes, while those for Ast_*Sfrp1* and Ast_*S100a4* were grouped as S100a11 trajectory marker genes. To determine the ‘on and off’ timing of expression of each gene, we used function geneSmoothFit() from URD, which takes a group of genes and cells, averages gene expression (parameters: moving.window = 5, cells.per.window = 25, pseudotime.per.window = 0.005), and then uses smoothing algorithms (spline fitting) to describe the expression of each gene^16^. Genes were then ordered and visualized according to their expression along the pseudotime via the function determine.timing() provided in URD package. The gene cascade heatmap was split into two (*i*.*e*., S100a11 and Olig2 trajectories) after the first branching point of the URD tree.

### PCA distance

PCA distance was calculated to measure the molecular changes of progenitor subtypes across ages (Extended Data Fig. 4e). Briefly, for each progenitor subtype (*i*.*e*., Prog_1 and Prog_2) at each timepoints (*i*.*e*., E10.5, E12.5, E14.5 and E16.5), the first 30 PCA dimensions were considered. The average PCA distance within for each progenitor subtype between E10.5 and the others timepoints was calculated using the “dist()” function from the R stats package. As a way of normalization, each average PCA distance between E10.5 and other ages was divided by a scrambled distance, which was calculated by randomly splitting the whole progenitor dataset into 2 groups of cells and calculating their average PCA distance.

## Data availability

scRNA-seq and MERFISH data will be deposited on GEO and be publicly available as of the date of publication.

## Code availability

This paper does not report original code. Codes and pipelines used in this study can be provided upon request.

## Acknowledgements

We thank the Genomics Platform, Bioimaging, and FACS Facility at the University of Geneva, as well as the MPIB NGS Platform for their invaluable support; A. Benoit for technical assistance; the entire Jabaudon laboratory their thoughtful feedback on the manuscript and for their constructive contributions throughout the project. The Bocchi laboratory and this project are supported by the Swiss National Science Foundation (Ambizione grant: PZ00P3_201995).

## Author contributions

R.B. conceived the project and designed experiments. J.Z. and R.B. performed experiments and analyzed data with help of all the authors. J.Z. and R.B. wrote the manuscript with input from all authors.

## Extended data figures

**Extended Data Fig. 1.**
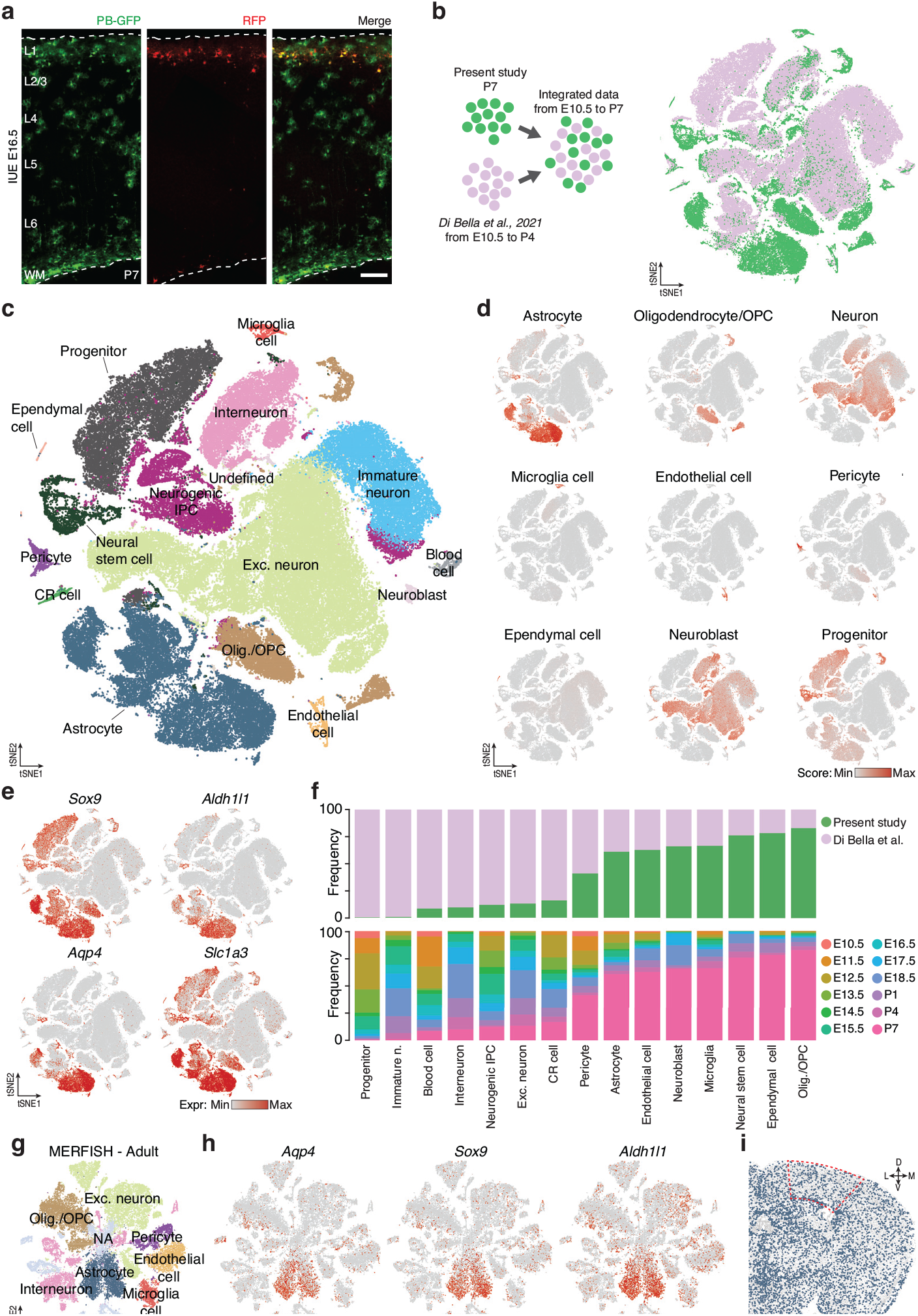
Integration and annotation of mouse embryonic and postnatal scRNA-seq and adult cortical MERFISH datasets. **a**, Representative confocal images of an entire cortical column in a P7 coronal section electroporated at E16.5 with both integrative (PB-GFP) and episomal (RFP) plasmids. Scale bar: 100 μm. IUE: *in utero* electroporation. **b**, tSNE plot illustrating the integrated dataset. Cells from present study depicted in green while those from Di Bella et al. are in pink. **c**, tSNE plot showing cell types in the integrated dataset. CR cell: Cajal–Retzius cell; IPC: intermediate progenitor cell; Olig./OPC: oligodendrocyte and oligodendrocyte precursor cell; Exc. Neuron: excitatory neuron. **d**, tSNE plots showing cell type scores, calculated based on known markers for major cell types in the cortex. **e**, tSNE plots showing the expression of four canonical astrocytic markers in the integrated dataset. **f**, Stacked bar graphs showing the proportion of cells from different studies (top) or from ages (bottom) for each cell type. **g**, tSNE plot showing cell types found in the adult MERFISH dataset. **h**, tSNE plots showing the expression of three canonical astrocytic markers in the adult MERFISH dataset. **i**, Spatial positioning in a coronal section of each individual cell of the adult MERFISH dataset. Astrocytes are highlighted in blue.

**Extended Data Fig. 2.**
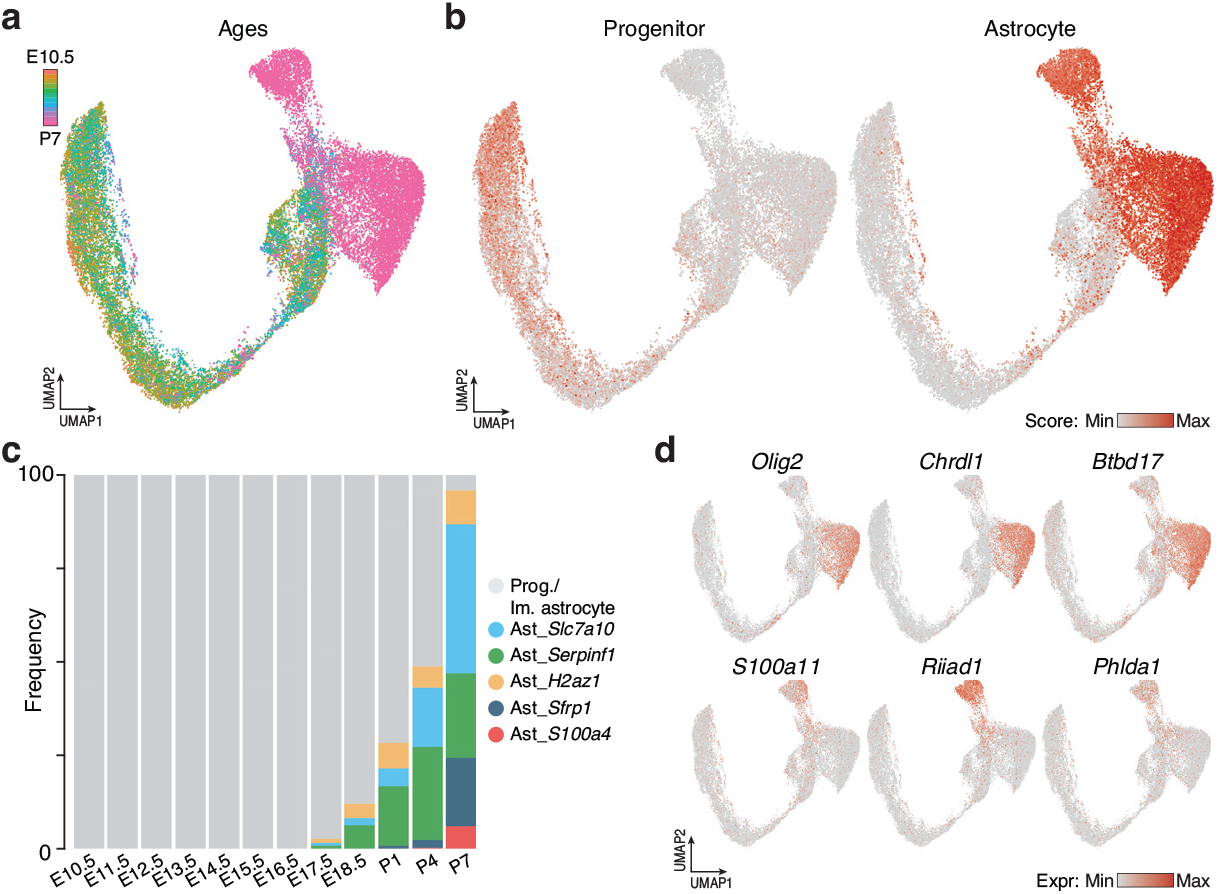
Characterization of cortical astrocyte trajectories. **a**, UMAP plot displaying progenitors and astrocytes aged from E10.5 to P7 color-coded according to their ages. **b**, UMAP plots showing progenitor and astrocyte scores. **c**, Stacked bar graph showing the relative proportion of the different astrocyte subtypes over time. Prog.: progenitor; Im. astrocyte: immature astrocyte. **d**, UMAP plots showing the expression of selected trajectory-specific genes.

**Extended Data Fig. 3.**
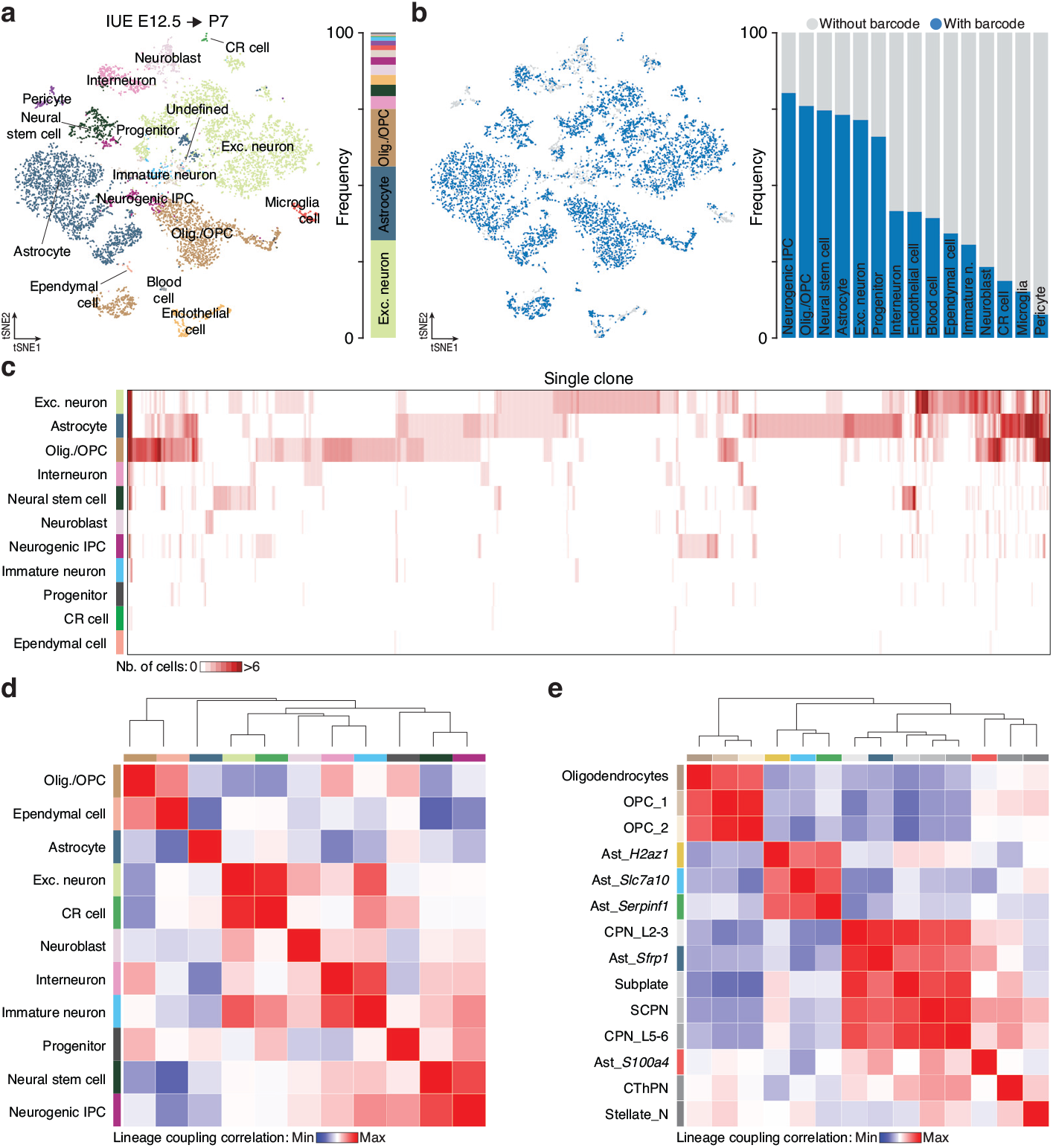
Lineage coupling analyses of major cell types in the mouse neocortex. **a**, Left: tSNE plot displaying all major cell types traced from E12.5 and collected at P7. Right: stacked bar plot showing the proportion of each cell type. **b**, tSNE plot highlighting cells tagged by TrackerSeq barcodes (left) with a histogram displaying the relative proportion of cells with and without barcodes in each major cell type (right). **c**, Heatmap showing clonal distributions across major cell types. Each column represents one clone, with number of cells per clone indicated by color scale. Blood cell, Microglia cell, Pericyte and Endothelial cell are excluded from the distribution analyses. **d**, Heatmap of lineage coupling scores between pairs of cell types included in (c). Values range from positive (red, coupled) to negative (blue, anti-coupled). **e**, Heatmap of lineage coupling scores between pairs of cell types further subdivided in subtypes (*e*.*g*., astrocyte subtypes, neuronal subtypes and OPC subtypes). Values range from positive (red, coupled) to negative (blue, anti-coupled).

**Extended Data Fig. 4.**
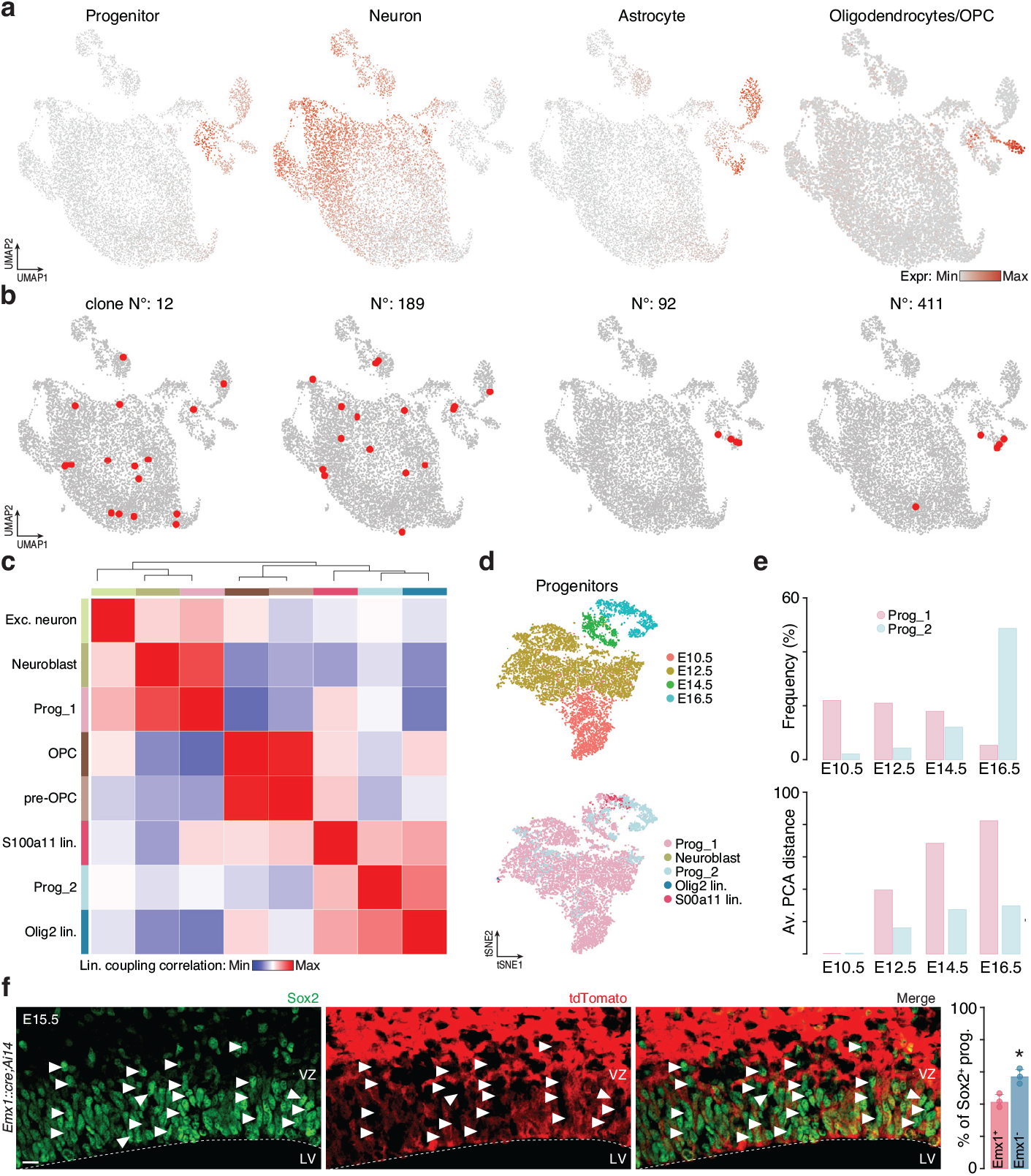
Identification and characterization of cortical progenitor subtypes. **a**, tSNE plots showing progenitor, neuron, astrocyte and Oligodendrocyte/OPC scores in the E18.5 dataset. **b**, Examples of clones that are either shared between neurons, Prog_1 and S100a11 lineage astrocytes (left) or restricted within Olig2 lineage astrocytes and Prog_2 (right). **c**, Heatmap of lineage coupling scores between pairs of major cell types identified in the E18.5 dataset. Values range from positive (red, coupled) to negative (blue, anti-coupled). **d**, tSNE plots showing progenitors from Di Bella et al. at E10.5, E12.5, E14.5 and E16.5, color-coded according to their ages (top) and by cell types predicted from the E18.5 dataset (bottom). **e**, Top: histogram showing the frequency distribution of progenitor subtypes across ages. Bottom: average PCA distance within each progenitor subtypes. The E10.5 time point was used as reference for each progenitor subtypes and the distance E10.5 to E12.5, E10.5 to E14.5 and E10.5 to E16.5 are represented. **f**, Representative confocal images of the in E15.5 VZ of a *Emx1::cre;Ai14* transgenic mouse immunostained by Sox2 labelling all progenitors. The proportion of tdTomato positive (Emx1^+^) and negative (Emx1^-^) cells are quantified on the right. Values are shown as mean ± s.d.; n = 3 experimental animals; two-tailed *t*-test, \**P* < 0.05. Emx1^-^ progenitors are indicated with white arrowheads. VZ: ventricular zone; LV: lateral ventricle. Scale bar: 20 μm.

**Extended Data Fig. 5.**
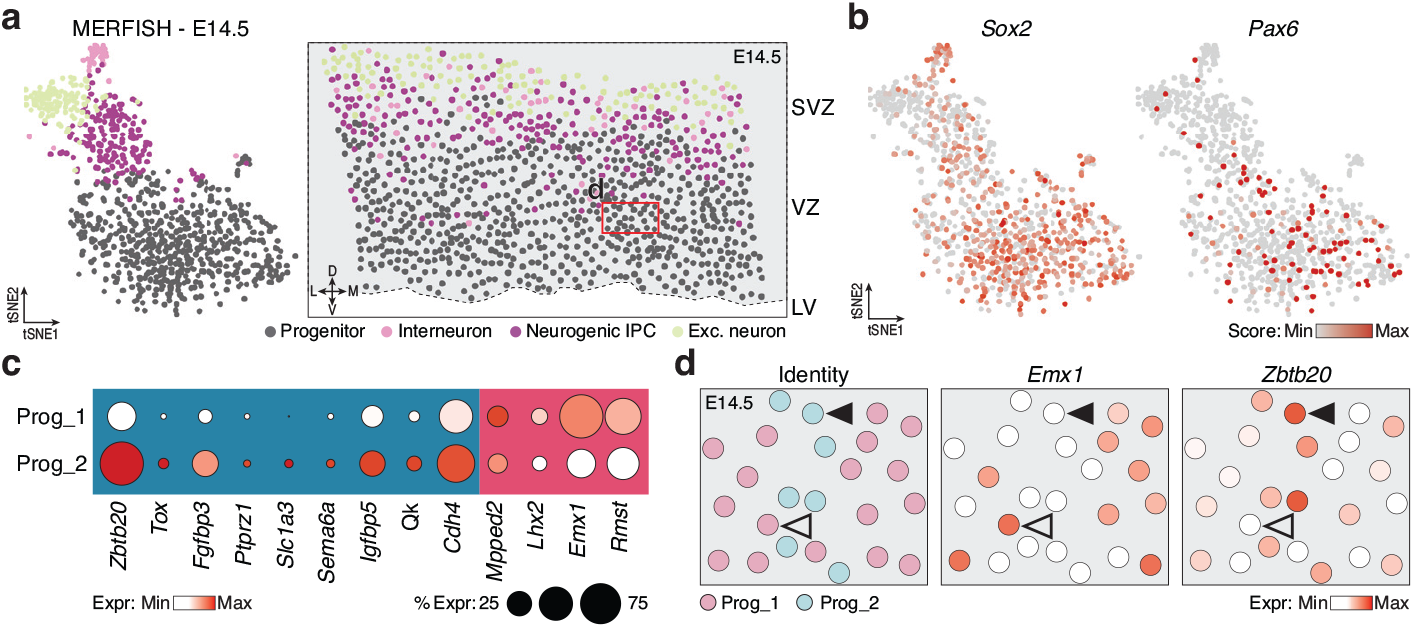
Spatial distribution of cortical progenitor subtypes. **a**, Left: tSNE plot showing cells from the E14.5 MERFISH dataset. Right: Spatial positioning in a coronal section of each individual cell of the E14.5 MERFISH dataset. **b**, tSNE plot showing the expression of two progenitor markers, *Sox2* and *Pax6*, in the E14.5 MERFISH dataset. **c**, Dot plot showing the expression level of selected genes for each progenitor subtypes. **d**, Expression of *Emx1* and *Zbtb20* in a cropped region of the coronal section of the E14.5 MERFISH dataset corresponding to the dashed region in panel **a**. Empty arrowheads indicate Prog_1 *Emx1*^*+*^*/ Zbtb20*^*-*^, while full arrowheads indicate Prog_2 *Emx1*^*-*^*/Zbtb20*^*+*^.

**Extended Data Fig. 6.**
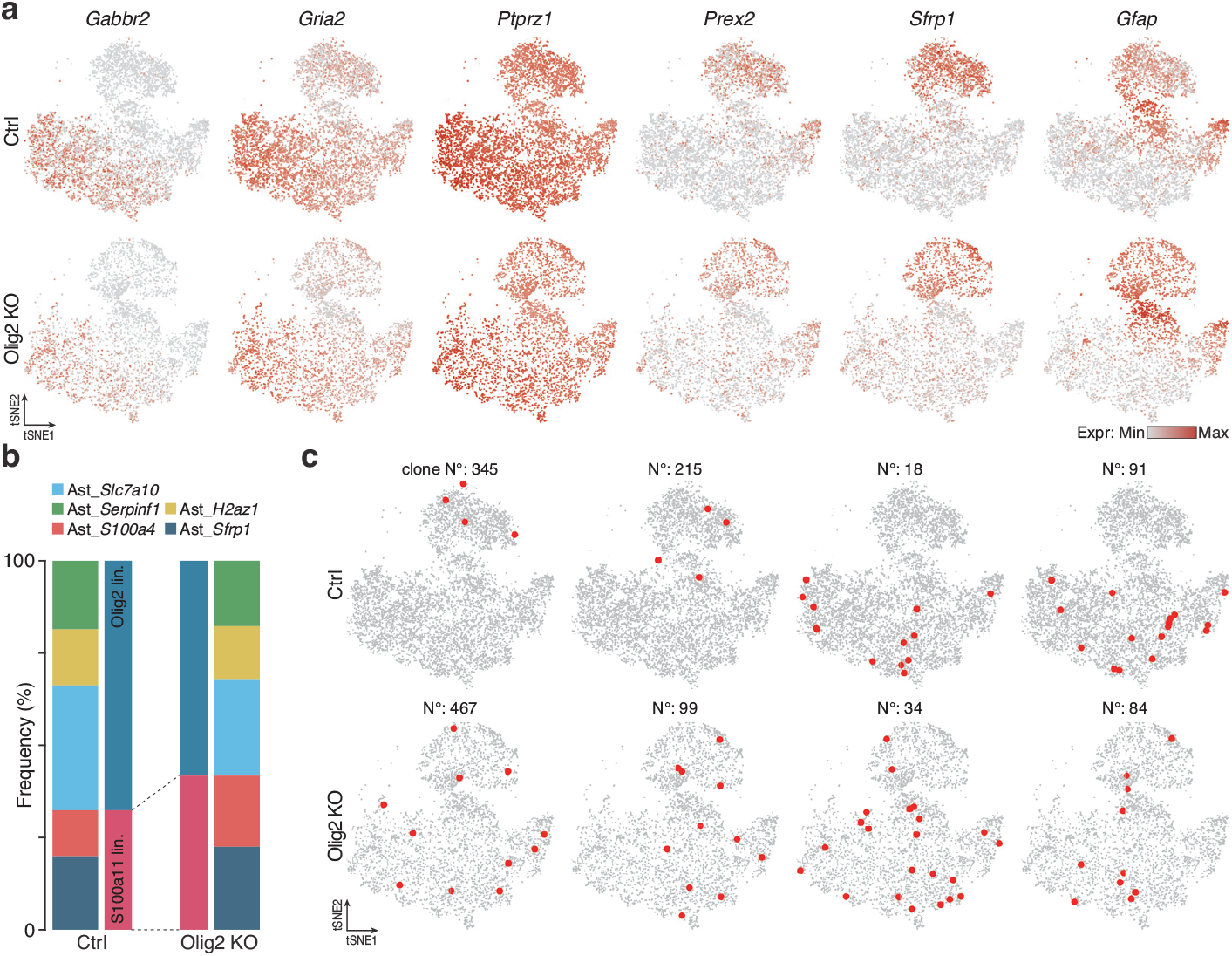
Alteration in transcriptome, subtype composition and clone distribution after *Olig2* knockout. **a**, tSNE plots showing the expression of three representative genes (*Gabbr2, Gria2* and *Ptprz1*) upregulated in Ctrl and three (*Prex2, Sfrp1* and *Gfap*) upregulated after Olig2 KO. **b**, Stacked bar plot showing the relative proportion of each astrocyte subtype in either Ctrl or Olig2 KO condition. **c**, Examples of clones in Ctrl or Olig2 KO condition. In Ctrl condition clones are mainly restricted within either Olig2 lineage astrocytes or S100a11 lineage astrocytes (top), whereas in Olig2 KO condition clones are shared between the two lineages (bottom).

